# Category-specific representational patterns in left inferior frontal and temporal cortex reflect similarities and differences in the sensorimotor and distributional properties of concepts

**DOI:** 10.1101/2021.09.03.458378

**Authors:** Francesca Carota, Nikolaus Kriegeskorte, Hamed Nili, Friedemann Pulvermüller

**Affiliations:** Max Planck Institute for Psycholinguistics, Wundtlaan 1, 6525 XD Nijmegen, The Netherlands; Donders Institute for Cognitive Neuroscience, RU, Kapittelweg 29, 6525, Nijmegen; Department of Experimental Psychology, University of Oxford, Tinbergen Building, 9 South Parks Road, Oxford OX1 3UD, UK; MRC Cognition and Brain Sciences Unit, University of Cambridge, Cambridge CB2 7EF, UK; Department of Psychology, Department of Neuroscience, and Department of Electrical Engineering, Zuckerman Mind Brain Behavior Institute, Columbia University, New York, NY 10032, USA; Brain Language Laboratory, Department of Philosophy and Humanities, WE4, Freie Universität Berlin, Berlin, Germany; School of Mind and Brain, Humboldt Universität zu Berlin, Berlin, Germany

**Author notes:** Corresponding author: Francesca Carota, Wundtlaan 1, 6525 XD Nijmegen, The Netherlands.

**Keywords:** sensorimotor semantics, symbol grounding, conceptual representations, Representational Similarity Analysis, fMRI

## Abstract

Neuronal populations code similar concepts by similar activity patterns across the human brain’s networks supporting language comprehension. However, it is unclear to what extent such meaning-to-symbol mapping reflects statistical distributions of symbol meanings in language use, as quantified by word co-occurrence frequencies, or, rather, experiential information thought to be necessary for grounding symbols in sensorimotor knowledge. Here we asked whether integrating distributional semantics with human judgments of grounded sensorimotor semantics better approximates the representational similarity of conceptual categories in the brain, as compared with each of these methods used separately. We examined the similarity structure of activation patterns elicited by action- and object-related concepts using multivariate representational similarity analysis (RSA) of fMRI data. The results suggested that a semantic vector integrating both sensorimotor and distributional information yields best category discrimination on the cognitive-linguistic level, and explains the corresponding activation patterns in left posterior inferior temporal cortex. In turn, semantic vectors based on detailed visual and motor information uncovered category-specific similarity patterns in fusiform and angular gyrus for object-related concepts, and in motor cortex, left inferior frontal cortex (BA 44), and supramarginal gyrus for action-related concepts.

## Introduction

Distributional statistics, capturing the meanings of words and concepts by the company they take, have been proposed to provide a basis for language comprehension (e.g., Landauer and Dumais, 1997; Landauer, 1998). Recent brain imaging research taking advantage of the information carried by population codes (e.g., neurons or voxels) in the brain has demonstrated that similar word meanings are represented by similar activity patterns (e.g., Kriegeskorte et al., 2008; Devereux et al., 2013; Carlson et al., 2014; Carota et al., 2017) and that the statistical distributions of words, as captured by co-occurrence frequency vectors (Landauer and Dumais, 1997; Mikolov et al., 2013), relate to such representational patterns in the language networks (e.g., Carlson et al., 2014; Pereira et al., 2018; Carota et al., 2021). However, how the abstract linguistic information that can be extracted from patterns of word co-occurrences relates to conceptual structure and its underlying neural representations remains unclear.

There is increasing agreement that distributional statistics in isolation fail to provide a cognitively realistic model of semantic knowledge (e.g., Barsalou, 2017), because it does not clarify the links between word symbols and ‘the world’ (referential semantics). To be meaningful, symbols must be anchored, or *grounded* (see Harnad, 1990; 2012; Gallese and Lakoff, 2005; Glenberg & Robertson, 2000; Barsalou, 2008; 2017; Marino et al., 2012; Pulvermüller, 2013), in the experiences of perceptions and actions that link them up with their extralinguistic referents - for example the object-related word *peach* with grounded semantic features, such as [+ROUND], [+YELLOW] and the action word *hike* with [+LEGS], [+MOVEMENT]. Such experiential attributes form an essential component of conceptual structure (Binder et al., 2016). They influence reaction times during semantic judgments and lexical decision tasks (Paivio, 1971; Barsalou, 1999; Barsalou et al., 2008; Vigliocco et al., 2009, 2014; Andrews et al., 2014; Kousta et al., 2011; Zwaan, 2014), and are reflected in the spatial localization (Hauk et al., 2004; Martin, 2007; Simmons et al., 2007; Fernandino et al., 2016) and temporal dynamics of brain activity (Pulvermüller et al., 2005; Moseley et al., 2013). Psycholinguistic theory holds that such experiential knowledge is linguistically encoded as a function of language use and task at hand (Symbol Interdependency hypothesis: Louwerse, 2008), so that an increase in perceptual simulation allows for richer and more detailed conceptual representations. Corroborating this view, recent behavioural results suggest that the sensorimotor properties of concepts enable a more precise specification of semantic categories, also when compared to distributional statistics *per se* (e.g.: Binder et al., 2016). Furthermore, when distributional and sensorimotor semantics are combined together, brain activation patterns associated to sentence meanings can be decoded with increased accuracy than using distributional information alone (Andrews et al., 2019).

Here, we investigate the relative contribution of distributional and sensorimotor information to the mapping of the brain’s multidimensional semantic space of concepts. We asked to what extent sensorimotor and distributional semantic information provide distinct and/or interdependent contributions to the neural encoding of conceptual meaning, and how the similarity structure in the corresponding neurocognitive representations relates to (and complements) to the similarity structure of the neural patterns of critical semantic regions in the language network.

It is well established that different sensory and motor attributes of object concepts relate to the brain activation patterns depending on how pertinent they are for defining that concept (Fernandino et al., 2016). For example, properties like colour (e.g., Simmons et al., 2007) and shape (e.g., Wheatley et al., 2005), differentiate object classes such as animals, tools and foods in inferior temporal and anterior ventrolateral frontal regions, whilst motion (e.g., Hauk et al., 2008), and sound (e.g., Kiefer et al., 2008) distinguish animals from foods in posterior middle and superior temporal cortex. In turn, taste and manipulation are distinctive properties of foods and tools, engaging orbitofrontal cortex (e.g., Barrós-Loscertales et al., 2012; Carota et al., 2012), and motor and posterior middle temporal cortex (e.g., Hoenig et al., 2008), respectively. Current theories make however different claims about how the different unimodal sensory and motor features of the meaning of a concept (for instance, for the object word *flute*, the shape, form, colour, sound etc. of typical referent objects) are integrated in multimodal regions in higher-order association cortices. According to the Hub-and-Spoke model, the anterior temporal cortex (aTL: Patterson et al., 2007) is an apex for all forms of semantic knowledge (i.e., both sensorimotor and distributional) to be brought together (Lambon-Ralph, 2017). A different model ascribes the role of a getaway to meaning to the middle temporal gyrus (MTG: Turken and Dronkers, 2011; Hickok, 2014), whilst another one emphasises the multimodal convergence of unimodal semantic features in inferior temporal gyrus (ITG, Price et al., 2000). Recent evidence supports the competing view that the angular gyrus in inferior parietal cortex (AG: Geschwind, 1965) is responsible of binding together unimodal information (e.g., Binder et al., 2009; Fernandino et al., 2016), whereas pars orbitalis (BA 47) of inferior frontal cortex (Poldrack et al., 1999) may enable semantic unification of linguistic units (e.g., Hagoort, 2005). The multimodal regions supporting the integration of unimodal features are also good candidates for interfacing sensorimotor and distributional information.

In particular, recent evidence suggested that the angular gyrus may be important to link distributional representations of concepts and their action-related properties, whereas anterior aspects left inferior frontal gyrus (LIFG, BA 45-47), important for semantic combinatorial operations unifying simpler meaningful units (e.g., words) into larger semantic structures (e.g., sentences) (e.g., Hagoort, 2005; 2016; 2019; Hagoort and Indefrey, 2014), may primarily support representational similarity based on the distributional statistics (Carota et al., 2020). The relevance of this region for semantic combinatorics has been explained as rooted in its executive – rather than representational – functions (e.g., Hoffman et al., 2018).

This earlier evidence leads to the competing hypothesis that sensorimotor semantics is predominantly represented in multimodal association regions, with only specific sensory and motor representations reaching modality-preferential cortex for specific semantic categories, whilst distributional semantics is primarily reflected in combinatorial semantic regions in frontal cortex.

Intriguingly though, some studies indicate that also modality-specific visual and motor regions may take a role in representing distributional similarities for specific semantic word categories (Mitchell et al., 2008; Carlson et al., 2014; Carota et al., 2017). For instance, distributional semantic similarities among words were indexed in posterior inferior temporal cortices (e.g., Carlson et al., 2014), known to support the processing of objects and for which previous studies demonstrated category selectivity for different types of nouns (e.g., places, foods, tools, and animals) (Damasio et al., 1996; Mitchell et al., 2008; Mitchell and Cusack, 2016). This leads to the competing hypothesis that these qualitatively distinct sources of semantic information may be redundantly represented both in higher-order association cortex and in lower-level category-preferential regions.

To test these hypotheses, we investigated to what extent a multidimensional vector space combining grounded semantic information (based on human ratings of experiential properties, such as colour, shape, sound, imageability, action-relatedness, taste, odour, arousal, and valence, etc.) and distributional statistics (modelled using GloVe: Pennington et al., 2014) captured the semantic similarity structure of word representations better than each of these models in isolation. We then asked to what extent cortical areas in higher-order association cortex (e.g., LIFG, aTL, MTG/ITG, AG) and in lower-level unimodal cortex (sensory and motor cortex) indexed the neurometabolic correlates of the semantic similarities between words and whether such representational similarity mapping differed spatially between different word types (action and object related words). We reanalyzed an existing dataset of recorded fMRI responses (Carota et al., 2021) while subjects silently read a highly controlled set of action- and object-related words and pressed a button when they detected an occasionally misspelled word. We related grounded and distributional properties of words to brain activity patterns by adopting Representational Similarity Analysis (RSA) (Kriegeskorte et al. 2008) to measures the co-variation of fMRI responses across multiple neighboring voxels within relevant brain areas of the semantic network.

In this context, the method helps to reveal the similarities and differences in the qualitative properties that inform semantic information encoding within regions of the language network that classical univariate activation-based approaches have shown to support either general multimodal semantic processes or category-specific processing of action and object word categories, by elucidating the relative contribution of distributional and sensorimotor information to the mapping of the brain’s multidimensional semantic space.

## Materials and Methods

### Participants

Twenty-three healthy volunteers participated in the study. All participants were right-handed (laterality quotient of 90, standard error (SE) = 3.1, Oldfield, 1971), monolingual English native speakers. Their mean age was 29 years (SE=2.8). Participant had no history of neurological or psychiatric disorders. They had normal or corrected-to-normal vision. All participants gave their informed consent to take part in the study and were remunerated for their time. Ethical approval was obtained from the Cambridge Psychology Research Ethics Committee.

### Experimental procedure

#### Stimuli

96 word stimuli, 16 from each individual category of leg-, arm-, face-related actions and tool-, animal-, food-related objects, were included in the study. Stimulus word groups were matched for word length (counted in number of letters), letter bigram and trigram frequency, logarithmic word frequency, number of orthographic neighbors, and standardized lexical frequency (see Behavioral Results). Relevant values were obtained from the CELEX database (Baayen et al. 1993) and the WordSmyth Web site (www.wordsmyth.net/). The word categories were selected based on differences in their rated semantic relationship to objects, actions, bodily sensations, emotional features, as well as their concreteness, and imageability (for discussion, see Pulvermüller 1999) (see Table 1). Strings of meaningless hash marks matched in length to the stimulus words were used as low-level baseline stimuli during 120 trials. Null events consisting of a fixation cross displayed at the centre of the screen were presented during additional 60 trials. 60 trials consisting of misspelled words to be detected by the participants throughout the experimental task were presented. These “typo” trials did not include words from any of the semantic categories from which the 96 target words were taken - so as to avoid a bias towards one of these categories - and were discarded from the analysis. Statistical comparisons were carried out between brain activation patterns elicited by matched word categories.

**Table 1.**
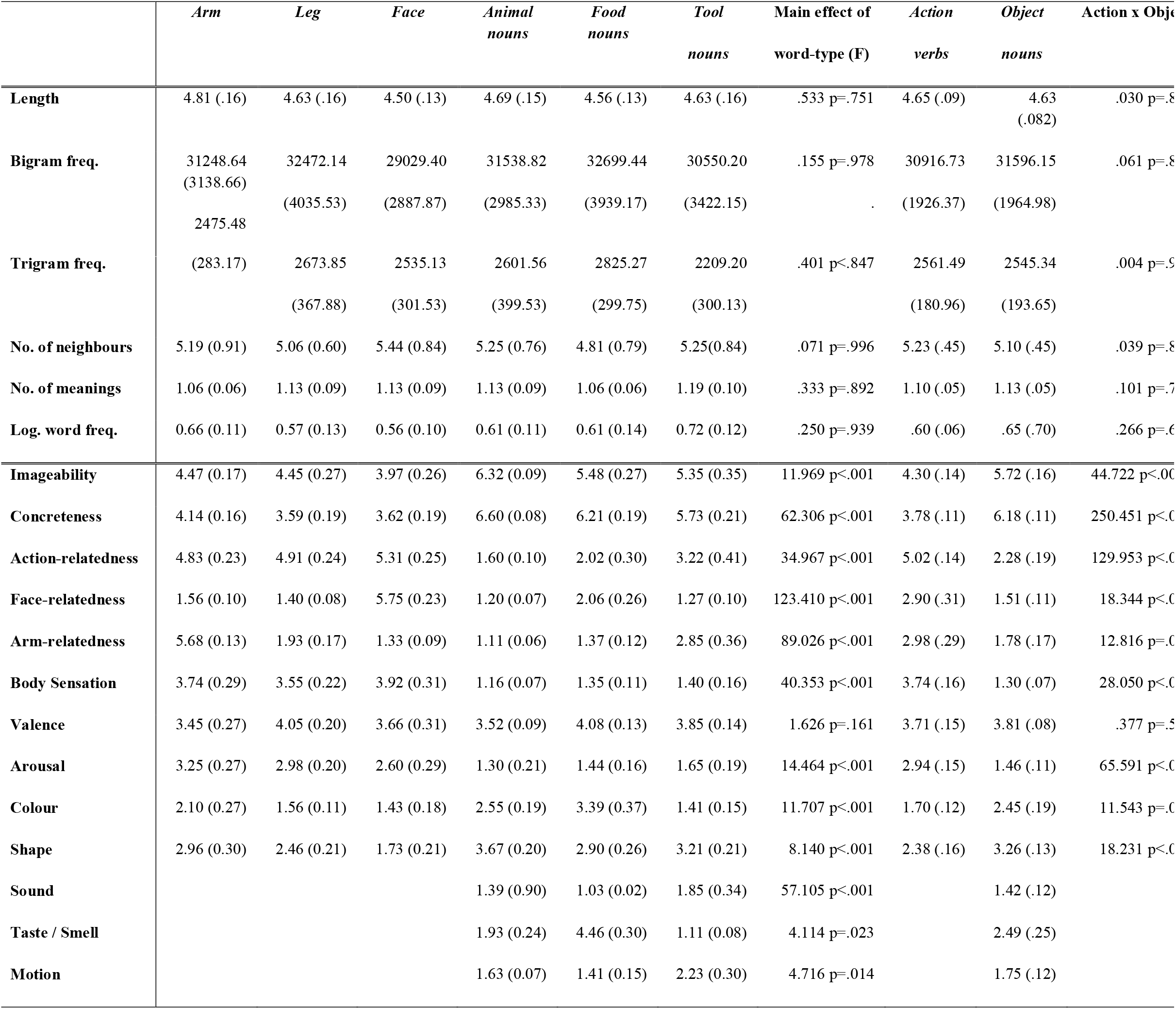
Psycholinguistic properties and semantic ratings are shown for each word sub-category, as well as for the categories of action- and object-related words. Means and standard errors (in brackets) are reported for each word category, along with results of an ANOVA comparing ratings between word groups.

|  | *Arm* | *Leg* | *Face* | *Animal*<br>*nouns* | *Food*<br>*nouns* | *Tool*<br>*nouns* | Main effect of<br>word-type (F) | *Action*<br>*verbs* | *Object*<br>*nouns* | Action x Object |
| --- | --- | --- | --- | --- | --- | --- | --- | --- | --- | --- |
| **Length** | 4.81 (.16) | 4.63 (.16) | 4.50 (.13) | 4.69 (.15) | 4.56 (.13) | 4.63 (.16) | .533 p=.751 | 4.65 (.09) | 4.63<br>(.082) | .030 p=.8 |
| **Bigram freq.** | 31248.64<br>(3138.66) | 32472.14<br>(4035.53) | 29029.40<br>(2887.87) | 31538.82<br>(2985.33) | 32699.44<br>(3939.17) | 30550.20<br>(3422.15) | .155 p=.978 | 30916.73<br>(1926.37) | 31596.15<br>(1964.98) | .061 p=.8 |
|  | 2475.48 |  |  |  |  |  | . |  |  |  |
| **Trigram freq.** | (283.17) | 2673.85 | 2535.13 | 2601.56 | 2825.27 | 2209.20 | .401 p<.847 | 2561.49 | 2545.34 | .004 p=.9 |
|  |  | (367.88) | (301.53) | (399.53) | (299.75) | (300.13) |  | (180.96) | (193.65) |  |
| **No. of neighbours** | 5.19 (0.91) | 5.06 (0.60) | 5.44 (0.84) | 5.25 (0.76) | 4.81 (0.79) | 5.25(0.84) | .071 p=.996 | 5.23 (.45) | 5.10 (.45) | .039 p=.8 |
| **No. of meanings** | 1.06 (0.06) | 1.13 (0.09) | 1.13 (0.09) | 1.13 (0.09) | 1.06 (0.06) | 1.19 (0.10) | .333 p=.892 | 1.10 (.05) | 1.13 (.05) | .101 p=.7 |
| **Log. word freq.** | 0.66 (0.11) | 0.57 (0.13) | 0.56 (0.10) | 0.61 (0.11) | 0.61 (0.14) | 0.72 (0.12) | .250 p=.939 | .60 (.06) | .65 (.70) | .266 p=.6 |
| **Imageability** | 4.47 (0.17) | 4.45 (0.27) | 3.97 (0.26) | 6.32 (0.09) | 5.48 (0.27) | 5.35 (0.35) | 11.969 p<.001 | 4.30 (.14) | 5.72 (.16) | 44.722 p<.001 |
| **Concreteness** | 4.14 (0.16) | 3.59 (0.19) | 3.62 (0.19) | 6.60 (0.08) | 6.21 (0.19) | 5.73 (0.21) | 62.306 p<.001 | 3.78 (.11) | 6.18 (.11) | 250.451 p<.001 |
| **Action-relatedness** | 4.83 (0.23) | 4.91 (0.24) | 5.31 (0.25) | 1.60 (0.10) | 2.02 (0.30) | 3.22 (0.41) | 34.967 p<.001 | 5.02 (.14) | 2.28 (.19) | 129.953 p<.001 |
| **Face-relatedness** | 1.56 (0.10) | 1.40 (0.08) | 5.75 (0.23) | 1.20 (0.07) | 2.06 (0.26) | 1.27 (0.10) | 123.410 p<.001 | 2.90 (.31) | 1.51 (.11) | 18.344 p<.001 |
| **Arm-relatedness** | 5.68 (0.13) | 1.93 (0.17) | 1.33 (0.09) | 1.11 (0.06) | 1.37 (0.12) | 2.85 (0.36) | 89.026 p<.001 | 2.98 (.29) | 1.78 (.17) | 12.816 p=.001 |
| **Body Sensation** | 3.74 (0.29) | 3.55 (0.22) | 3.92 (0.31) | 1.16 (0.07) | 1.35 (0.11) | 1.40 (0.16) | 40.353 p<.001 | 3.74 (.16) | 1.30 (.07) | 28.050 p<.001 |
| **Valence** | 3.45 (0.27) | 4.05 (0.20) | 3.66 (0.31) | 3.52 (0.09) | 4.08 (0.13) | 3.85 (0.14) | 1.626 p=.161 | 3.71 (.15) | 3.81 (.08) | .377 p=.5 |
| **Arousal** | 3.25 (0.27) | 2.98 (0.20) | 2.60 (0.29) | 1.30 (0.21) | 1.44 (0.16) | 1.65 (0.19) | 14.464 p<.001 | 2.94 (.15) | 1.46 (.11) | 65.591 p<.001 |
| **Colour** | 2.10 (0.27) | 1.56 (0.11) | 1.43 (0.18) | 2.55 (0.19) | 3.39 (0.37) | 1.41 (0.15) | 11.707 p<.001 | 1.70 (.12) | 2.45 (.19) | 11.543 p=.001 |
| **Shape** | 2.96 (0.30) | 2.46 (0.21) | 1.73 (0.21) | 3.67 (0.20) | 2.90 (0.26) | 3.21 (0.21) | 8.140 p<.001 | 2.38 (.16) | 3.26 (.13) | 18.231 p<.001 |
| **Sound** |  |  |  | 1.39 (0.90) | 1.03 (0.02) | 1.85 (0.34) | 57.105 p<.001 |  | 1.42 (.12) |  |
| **Taste / Smell** |  |  |  | 1.93 (0.24) | 4.46 (0.30) | 1.11 (0.08) | 4.114 p=.023 |  | 2.49 (.25) |  |
| **Motion** |  |  |  | 1.63 (0.07) | 1.41 (0.15) | 2.23 (0.30) | 4.716 p=.014 |  | 1.75 (.12) |  |

After the fMRI experiment, participants completed an unannounced word recognition test containing both novel distractor and experimental words. They performed above chance (average hit rate: 85% [STD: 8.3%]), indicating their attention to the words and compliance with the task. Results were used to confirm that subjects had been attentive continuously during the silent reading task.

#### Experimental Design

We adopted a rapid, periodic single trial, event-related paradigm. Fixation cross was presented at the centre of the screen whenever no stimulus was shown. Word and hash string stimuli were presented tachistoscopically, to discourage subjects from eye movements, for 100 ms. The stimulus onset asynchrony (SOA) was randomly varied between 3.5 and 4s. This design yielded overlapping, yet detectable hemodynamic responses (Kriegeskorte et al. 2008; Nili et al. 2014). The 96 stimulus words were presented in a different pseudo-random order in each of the 4 runs. Each stimulus occurred once per run (4 repetitions of each word in total). Stimuli were visually presented by means of E-Prime software (Psychology software Tools, Inc., Sharpsburg, USA, 2001) through a back-projection screen positioned in front of the scanner and viewed on a mirror placed on the head coil.

#### Task

Participants were engaged in a typo-detection task. They were given the instruction to attend to all the stimuli, to silently read the words and to understand their meaning. In addition, they were instructed to press a button with their left index finger if a misspelled word appeared at the centre of the screen.

### Imaging Methods

Subjects were scanned in a Siemens 3T Tim Trio using a head coil. Echo-planar imaging (EPI) sequence parameters were TR = 2000 msec, TE = 30msec, and flip angle = 78 degrees. The functional images consisted of 32 slices covering the whole brain (slice thickness 3mm, in-plane resolution 3 x 3mm, inter-slice distance 0.75mm).

### Data Analysis

Imaging data were analysed using SPM12 software (Wellcome Department of Imaging Neuroscience, London, UK). Images were corrected for slice timing and re-aligned to the first image using sinc interpolation. The EPI images were co-registered to the structural T1 images using a mutual coregistration procedure (Maes et al. 1997). The structural MRI was normalised to the 152-subject T1 template of the Montreal Neurological Institute (MNI). The resulting transformation parameters were applied to the co-registered EPI images. During the spatial normalization, images were resampled with a spatial resolution of 2 mm × 2 mm × 2 mm.

### Representational Similarity Analysis

For multivariate RSA (Kriegeskorte et al., 2008; Nili et al., 2014), the analysis was carried out in participant native space, using realigned, unsmoothed and non-normalised functional data, which were co-registered with MPRAGE of each subject. Data were analyzed using the general linear model. Response-amplitude was estimated for each voxel and for each of the 96 stimuli by performing single univariate linear model fit. Runs were concatenated along the temporal dimension. A separate hemodynamic predictor was included for each of the 96 stimulus words. As each word stimulus occurred once in each run, each of the 96 stimulus words had a distinct hemodynamic response per run, extended across all within-session runs. The time course of the predictors was determined based on the event sequence and a linear model of the hemodynamic response (Boynton et al., 1996). For each run, the design matrix was composed of the stimulus-response predictors with six head motion parameter time courses and a confound-mean predictor. The response-amplitude (beta) estimate map associated to each stimulus was converted into a t map by contrasting them against the implicit baseline in order to compute the representational dissimilarity matrices for RSA (Nili et al., 2014) (SI Materials and Methods).

### Model Specification

In order to test for the contribution of sensorimotor and distributional semantics on the representation of different word categories, we built up a semantic vector combining these qualitatively different properties, which were respectively assessed by semantic ratings and a corpus-based approach (See Fig. 1).

**Fig. 1.**
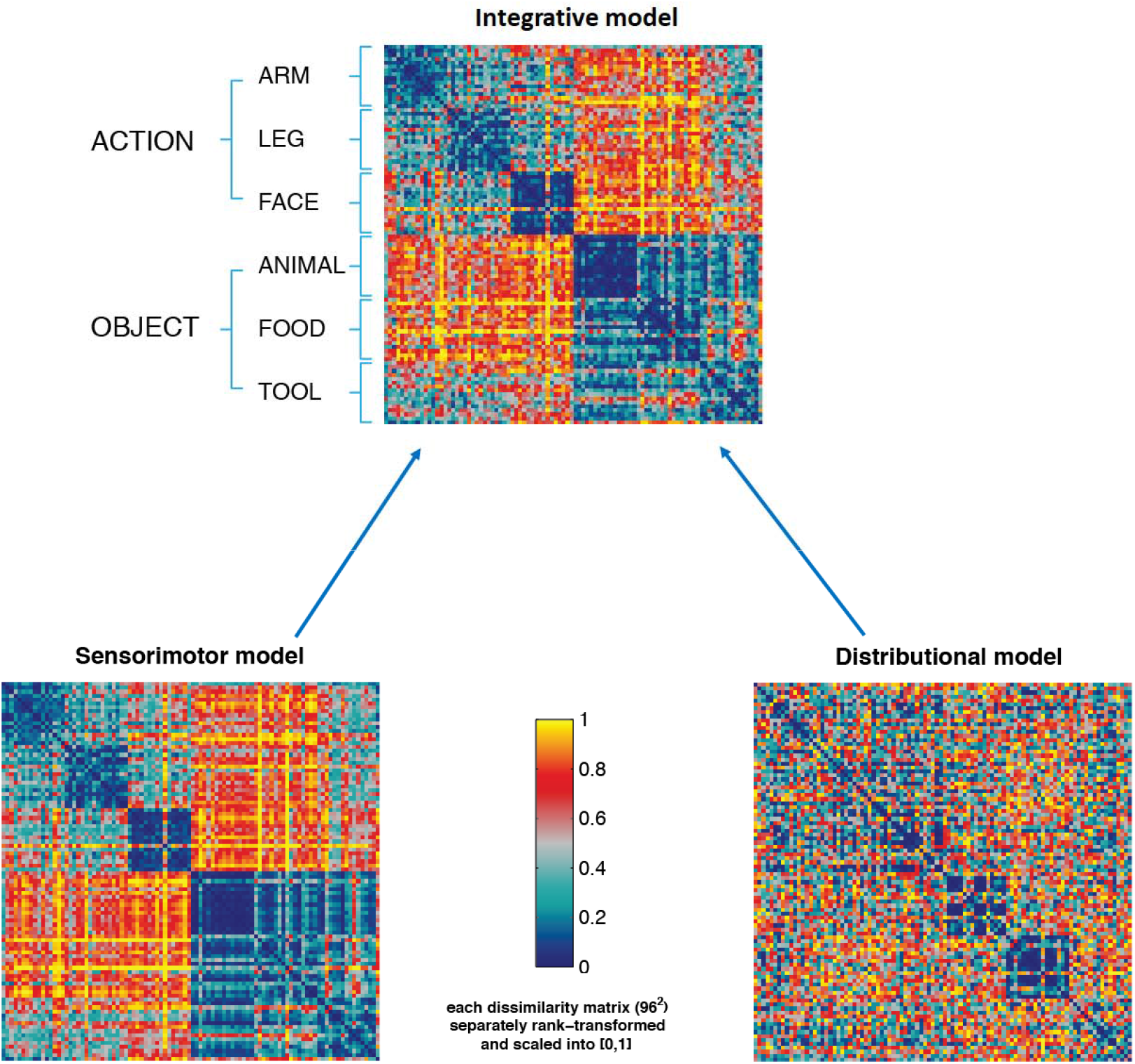
Representational dissimilarity matrices (RDM) displaying the semantic distances among the 96 stimulus words based on a) the grounded ‘sensorimotor’ model reflecting human ratings (bottom left); b) the distributional model based on GloVe (bottom right) and c) the integrative model (top panel). Dark blue (= 0) indicates maximal similarity between identity pairs and yellow (=1) indexes maximal dissimilarity. Each square in the model RDM shows (dis)similarities between the individual word items, which are plotted from top to bottom and from left to right – with dark blue indicating great semantic similarity or identity and red/yellow indexing dissimilarity. Therefore, the first diagonal in the matrix indicates the semantic identity of each single item with itself. Note that the distributional model does not reflect the subdivision of the experimental vocabulary into six semantic sub-categories (face, arm and leg related action verbs and animal, food and tool nouns), although the similarities within the categories of animal and especially that of food words differ from the background, appearing as blue rectangles of great semantic similarity. The sensorimotor model also shows the different noun categories, although the verb subcategories are not clearly delineated against each other by either of these models. Nevertheless, the large lexical and semantic category of action verbs is highlighted by a big blue square on the upper left of the left bottom panel. By far the best mapping of the categorical semantic structure of the vocabularies is achieved by the integrated model at the top (see Table 2 for statistical tests); please note the series of six blue squares form top left to bottom right representing semantic similarities within each of the six semantic subtypes. Also, the lexical categories of verbs and nouns are clearly revealed by this integrated model (big red/yellow ‘dissimilarity squares’ at lower left and upper right). The only exception from the between-lexical category-dissimilarity is seen for tool words, which are indeed semantically associated to action verbs (for example, “knife” and “carve”/ “peel”). We conclude that the integrated model is superior to the single method models in mapping semantic along with lexicosemantic similarities between the words used in this experiment.

#### Sensorimotor Model

The grounded ‘sensorimotor’ semantic model was constructed based on a number of action-related, visual and affective-emotional semantic properties relevant for the test words (for discussion, see Pulvermüller, 1999; and Binder et al., 2016), as rated in an independent study following established procedure (Pulvermüller, 1999; Hauk et al., 2008; Carota et al., 2017). These included motor properties, such as 1) action-relatedness (including, for objects such as foods and tools, association with edibility and manipulability: Carota et al., 2012), 2) body-part relatedness (e.g., arm-, leg- and face-relatedness), perceptual properties such as, 3) colour, 4) shape, 5) imageability, 6) motion 7) taste/smell, 8) sound and emotional properties, such as 9) arousal, and 10) valence. Semantic similarity between each pair of words was measured as the correlation distance (1-r) between the values of the properties specific to each pair of words. The resulting correlation values were entered in each cell of a Representational Dissimilarity Matrix (RDM) (Fig. 1, left bottom panel). For the separate analyses of category-specific patterns, the sensorimotor model for action words reflected the properties 1-12, whilst properties 1-15 contributed to the model specific to object words.

#### Distributional Model

This model coded for the semantic similarity between words, i.e. the probability for each pair of words to occur in *similar* contexts, by applying a state-of-art extension of Latent Semantic Analysis (LSA, Landauer and Dumais, 1997), GloVe (Pennington et al., 2014) to the British National corpus (BNC). The BNC contains 4096 texts with samples of written and spoken English from a wide range of sources and a wide variety of genres for a total of 100 million words. The words of each text were lemmatized by grouping together different inflected forms of a same word and used to construct a matrix in which each textual fragment containing a given word was represented as a row, each lemma was represented as a column, and each cell expressed the frequency with which each word occurs with all the other words in the corpus. After transformation of these word frequency values in their logarithm, Singular Value Decomposition (SVD) was applied to this matrix. SVD mapped the word vector space to a lower dimensional vector space, effectively grouping similar contexts together. Semantic similarity between each pair of words was then measured as the cosine between two word-vectors: the smallest the cosine, the greatest the similarity between word stimuli pairs. The values of latent semantic distances were expressed as a dissimilarity matrix (Fig. 1, right bottom panel).

#### Integrative (Sensorimotor and Distributional) Model

A third semantic model integrating rated sensorimotor and affective properties and distributional properties of words, was obtained by averaging the values contained in the two model RDMs. The resulting RDM is displayed in the right panel of Fig. 1 (top panel). The integrative semantic model expressed the contribution of the distributional and sensorimotor information about the test words in terms of pairwise similarity and dissimilarity relations, positing a mutual relationship between the experiential properties of two words and their co-occurrence likelihood. In particular, the smaller the dissimilarity between the set of experiential properties for two items (e.g., for the pair peach / plum: similarity in shape, ‘round’; action-related, ‘edible/manipulable’, taste, odour, etc.), the higher their likelihood to co-occur in a similar and coherent semantic context (e.g., groceries at the market, or cooking recipe). Reversely, the bigger the experiential property dissimilarity between two items (e.g., peach and jive), the smaller their likelihood to fit similar contexts.

### Whole-brain RSA Searchlights

Data were extracted for each participant individually using a “sphere of information” searchlight approach (Kriegeskorte et al., 2008; Nili et al., 2014). A roaming spherical searchlight with 10mm radius (Kriegeskorte et al., 2008) was moved throughout the grey matter to extract continuous, voxel-by-voxel maps of word-elicited activation values. To achieve maximal sensitivity to our experimental manipulations, this analysis was based on single items, with each experimental word modeled as a condition and associated with a separate hemodynamic predictor. The correlation distances (1-Pearson’s correlation) between the response patterns for each word paired with every other word were expressed as representational dissimilarity matrices (RDMs), which are symmetric about a diagonal of zeros (Kriegeskorte et al., 2008). These brain data RDMs were then correlated with theoretical model RDMs (using Spearman’s rank correlation) at each brain location. The resulting maps of r values for each participant and model were normalised onto the MNI template and entered into a group-level random-effects (RFX) analysis using permutation-based non-parametric statistics in SNPM (http://www2.warwick.ac.uk/fac/sci/statistics/staff/academic-research/nichols/software/snpm), to test for positive correlations between the model RDMs and brain data RDMs and thus determine the brain regions where the models best explained the brain activity. FDR correction at 0.05 for multiple comparisons across voxels and number of models was applied. 10,000 permutations were used in the analysis. In order to ensure that the searchlight maps we reported did not suffer from distortion due to either the searchlight size or the detection of fewer informative voxels, we performed additional analyses testing for the relatedness of the models and the brain activity patterns in selected regions of interest, as described below.

### Analyses per regions of interest

#### Definition of ROIs

An exploratory analysis was run in pre-selected Regions of Interest (ROIs) in the left hemisphere with well-studied semantic roles in language comprehension (for reviews: Binder and Desai, 2011; Pulvermüller et al., 2013). These included the 1) LIFG (BA 44, BA 45 and BA47) (Poldrack et al. 1999; Hagoort, 2005); 2) superior, middle, and inferior temporal gyrus (STG, MTG, ITG, further divided into anterior (aSTG, aMTG, aITG) and posterior (pSTG, pMTG, pITG) sections: Hickok and Poeppel, 2007; Turken and Dronkers, 2011; Hickok, 2014; Price et al., 2000); 3) temporal pole, TP (Patterson et al., 2007; Lambon-Ralph, 2017); 4) fusiform gyrus (FuG: Mion et al., 2010); 5) precentral gyrus (PCG: Hauk et al., 2004); and 6) inferior parietal cortex (both supramarginal gyrus, SMG and angular gyrus, AG: Binder et al., 2009; Fernandino et al., 2016) (see Fig. 3.A). We automatically defined these ROIs using the standard Wake Forest University (WFU) Pickatlas toolbox, which generates ROI masks in standard MNI space based on the Automated Anatomical Labelling (AAL) parcellation. In order to carry out multivariate analysis within individual-subject native space, all ROI-masks were transformed to subject native space by inverting the spatial normalisation applied during GLM analysis.

**Fig. 2.**
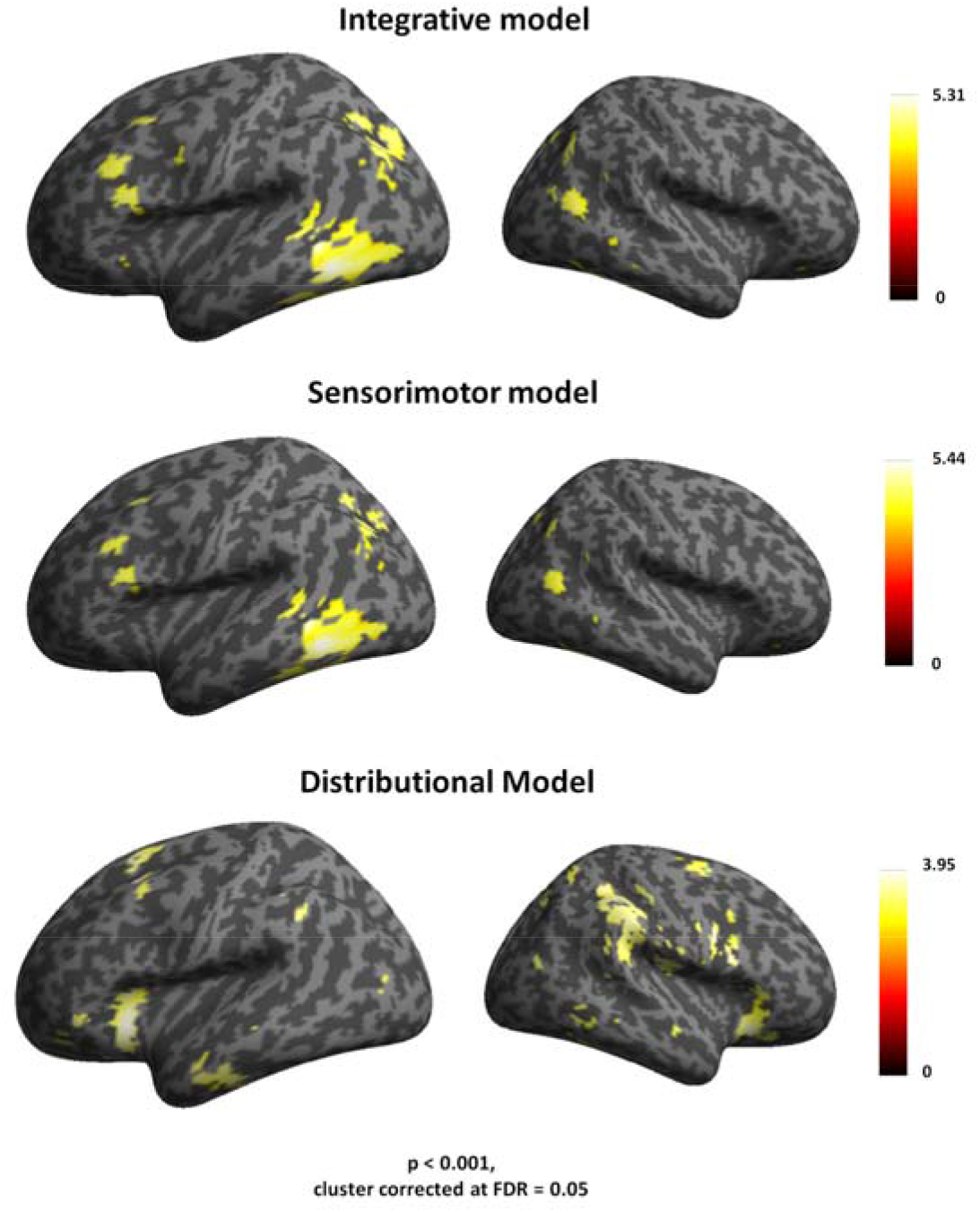
Results from whole-brain searchlight RSA. Top panel: the correlational effects of the grounded sensorimotor model in the pars opercularis of the inferior frontal cortex (BA 44), middle frontal cortex, inferior temporal and parietal cortex. Middle panel: the effects of the distributional LSA model emphasising the representational similarities based on co-occurrence information in higher-order regions in left inferior frontal cortex (BA 47) and anterior temporal cortex. Bottom panel: widespread activity triggered by the integrative model in distributed prefrontal, premotor, temporal and parieto-occipital cortex.

**Fig. 3.**
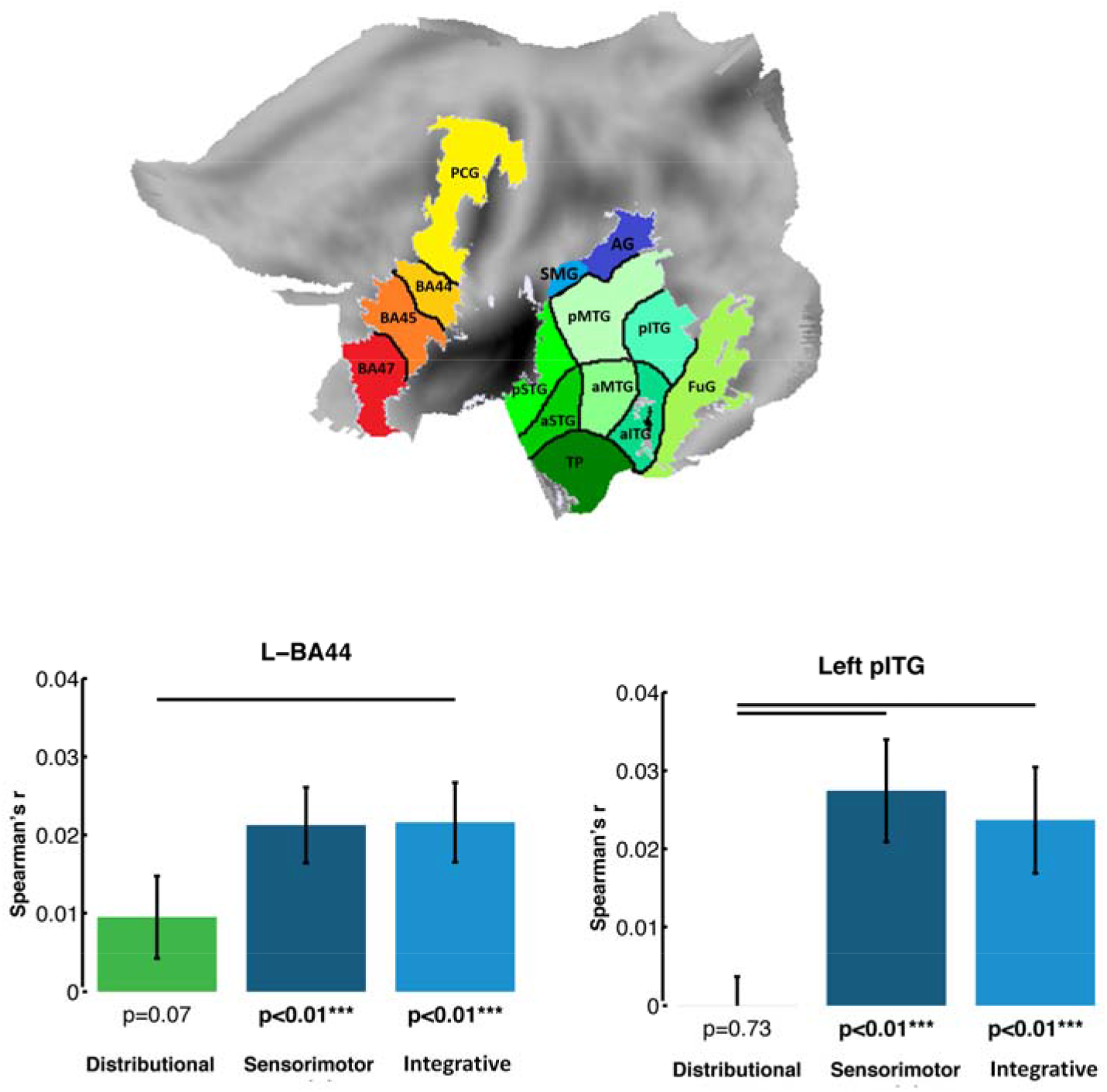
Results from model comparisons in selected ROIs, as displayed in the top section **A**. The bar graph depicts the averaged model-fMRI pattern correlations for each of the semantic model under examination (bottom section **B**). In each panel, the green bar indicates the distributional model, the blue bar the sensorimotor model and the dark blue bar the integrative model. Spearman’s rank correlations were calculated to assess the relatedness between brain activity and model RDMs and statistical inference was applied on the single subject correlations using a one-sided signed-rank test across subjects, testing whether the resulting correlation coefficients were significantly greater than zero. Below each bar, the significance value for the test is reported, corrected for multiple testing across brain regions by applying the FDR procedure; the expected FDR was less than 5% (Benjamini and Hochberg 1995). The horizontal bars in black indicate significant differences from model comparisons after FDR correction across models (FDR=0.05). Left panel: Effects specific to the LIFG (BA 44), where the distributional model differed significantly from both the sensorimotor and integrative models. Left panel: Effects specific to the left pITG. There was a significant difference (FDR=0.05) between the effects of the sensorimotor and integrative models and the ones triggered by the distributional model.

### Data cross-validation

To further estimate predictive performance and adjudicate between multiple models, ROI results were cross-validated using LDC (cross-validated Mahalanobis distance) on split data (Walther et al., 2015). In this approach, for each of k folds, k-1 of k independent subsets of the data (training set) are used to fit the parameters of each model and the left-out subset (test-set) is used to estimate predictive performance. Cross-validated LDC distances provide almost unbiased and conservative estimates of pattern dissimilarity. FDR correction at 0.05 for multiple comparisons across voxels and number of models was applied. 10,000 permutations were used in the analysis.

### Relatedness between fMRI patterns and models

RDMs were computed for each participant for the abovementioned ROIs. In each subject, the brain activity RDMs computed for each ROI were compared to model RDMs by calculating the Spearman’s rank correlation. To assess the relatedness between brain-activity and model RDMs, statistical inference was applied on the single-subject correlations using a one-sided signed-rank test across subjects, testing whether the resulting correlation coefficients were significantly greater than zero (Wilcoxon, 1945). We controlled the false discovery rate (FDR) for multiple comparisons of different brain regions by ensuring that the expected FDR was less than 5% (Benjamini and Hochberg, 1995). In order to test for the sensitivity of the model to semantic word categories, a first analysis in the whole brain (searchlights) and in selected ROIs was performed using the 96×96 matrix containing all words. A second ROI analysis was separately performed on the 48×48 sub-matrices containing the action words and the object words. Furthermore, these analyses first assessed the specific correlation of the distributional (LSA) and sensorimotor semantic models and brain activity patterns separately, and secondly the joint contribution of the two semantic dimensions here under focus. We directly tested the hypothesis that action and object words are respectively encoded in fronto-central and temporal regions which are engaged by the processing of grounded action-related and visual properties of words. To this aim, an ANOVA contrasted correlation values for action and object words in a hypothesis-driven sub-set of frontal and temporal regions associated with the processing of action and object word (Hauk et al., 2004). These ROIs covered 1) the pars triangularis (BA 45) and 2) and opercularis (BA 44) of the LIFG, 3) a premotor region which was non adjacent to inferior premotor cortex and BA 44 4) anterior temporal cortex (Patterson et al., 2007), 5) fusiform gyrus (Mion et al., 2014), and 6) angular gyrus (Binder and Desai, 2011). We additionally performed a 2×2 ANOVA (design: Region (6 levels, with 3 frontocentral and 3 temporo-parietal ROIs) x Word Category (2 levels: action and object words) to test for a double dissociation for the representations of action- and object-related words, contrasting r values words in the ROIs between word groups.

Furthermore, we analysed the discriminability of semantic word categories in the patterns of each ROI by comparing (t-test) the within-category similarity values for each semantic type with the between-category similarity values.

### Statistical model comparisons

Analyses were run to statistically compare the model-fMRI pattern fit in the abovementioned ROIs. A first analysis using the 96×96 matrix containing all words compared the performance of the distributional and sensorimotor models, as well as their integrative model, to assess their relative contribution to the mapping of semantic similarity in the brain’s semantic space. A second analysis testing for the sensitivity of the model to semantic word categories was separately performed on the 48×48 sub-matrices containing the action words and the object words. This analysis interrogated the sub-spaces of action and object words to test for differentiations in the underlying neural representations for distributional, sensorimotor, and integrative similarity. In particular, we tested the hypothesis that, if sensorimotor semantics - but not distributional statistics - is required for specific semantic representations (see Introduction), the corresponding similarity patterns would reflect distinctions for actions and object words only when their specific rated properties are activated. We therefore first compared the sensorimotor sub-models specific to action and object words. We then focused on dominant attributes of action and object words, known to be reflected in the corresponding action and perception systems. To this aim, two models were constructed, a first one coding for the rated motor properties (obtained from action-relatedness and leg-, arm-, face-relatedness) and a second one coding for the rated visual properties (colour, shape, imageability).

This was done by computing, for each subject, non-parametric Spearman□s rank correlations between model and brain activity RDMs. The two models were compared by subtracting the r-value of the correlation between the second model and the fMRI RDM from the r-value of the correlation between the first model and the fMRI RDM. The difference in r-value across all subjects was then tested against the null hypothesis of the value 0, to test for a difference in correlation, using a 2-sided Wilcoxon signed-rank test. P-values surviving FDR correction for multiple comparisons are reported (Benjamini and Hochberg 1995).

## Results

### Behavioural results

To evaluate the relationship between sensorimotor semantic properties and the six semantic word categories, results of the semantic rating experiment were evaluated. Mean values and standard errors for both semantic properties, for which word groups differed, and psycholinguistic variables, for which they were matched, are summarized in Table 1.

### Semantic similarities revealed by three linguistic models

The discriminability of semantic word categories in each model RDM was analysed by comparing (t-test) the within-category similarity values for each semantic type with the between-category similarity values. The resulting t-values were larger for the combination of sensorimotor ratings and distributional semantics (t= 6.30; p <0.0001) than for each individual model of sensorimotor information (t = 5.40; p<0.0001) and distributional semantics (t = 4.50; p<0.0001). This suggests that the combined model of integrative sensorimotor and distributional information produces better semantic category discrimination than each of the two measures alone, already on the level of purely cognitive-linguistic classification.

### Results from whole-brain RSA searchlights

Whole-brain RSA searchlights revealed that the integrative model correlated with the similarity structure of the neural patterns in fronto-temporal and parieto-occipital cortex (see Table 2 and the top panel of Fig. 2). A major cluster of activity was identified in left pITG and FuG, followed by a second one in left occipital lobe, and AG. A third cluster was seen in the pars orbitalis (BA 47), pars triangularis (BA 45) and pars opercularis of the LIFG (BA 44), extending to premotor cortex. Additional significant correlations were found in the right MTG, and, to a lesser extent, in parieto-occipital regions. The grounded sensorimotor model related to the neural patterns in the same regions as the integrative model, except the LIFG BA 47 (see Table 3 and Fig. 2, middle panel). Semantic similarity mapping of the distributional model led to pronounced correlations with the neural pattern in more anterior fronto-temporal regions, including bilateral IFG (BA 47), bilateral dorsal pre-supplementary motor area (pre-SMA), and AG, which showed pronounced correlations with the model in the right hemisphere (see Table 4 and bottom panel of Fig. 2).

**Table 2.** Results from searchlight RSA for the integrated model. Table of coordinates and significance voxel-level peak values (p) in each activation cluster that was correlated with the integrated semantic model integrating distributional and sensorimotor information.

| Regions | Cluster Extent | Voxel-level P | Pseudo T | Coordinates |  |  |
| --- | --- | --- | --- | --- | --- | --- |
|  |  |  |  | x | y | z |
| Left fusiform | 2850 | 0.017 | 4.2 | -39 | -46 | -24 |
| Left Inferior Temporal |  | 0.017 | 4.45 | -51 | -46 | -12 |
| Left Mid Occipital |  | 0.017 | 4.31 | -24 | -64 | -36 |
| Left Mid Frontal | 1021 | 0.017 | 4.68 | -33 | 23 | 48 |
| Left Insula |  | 0.018 | 4.14 | -30 | 23 | -1 |
| Left Inferior Frontal Opercularis |  | 0.026 | 3.77 | -51 | 11 | 29 |
| Left Precentral Gyrus | 120 | 0.030 | 3.60 | -45 | 11 | 32 |
| Right Middle Temporal | 1514 | 0.017 | 4.52 | 51 | -64 | 10 |
| Right Middle Temporal |  | 0.018 | 4.23 | 45 | -73 | 21 |
| Right Angular |  | 0.018 | 4.20 | 36 | -70 | 44 |
| Right Inferior Frontal Orbitalis | 1604 | 0.019 | 4.27 | 33 | 32 | -12 |
| Right Inferior Frontal Opercularis |  | 0.019 | 4.02 | 51 | 11 | 25 |
| Right Inferior Frontal Triangularis |  | 0.019 | 3.74 | 48 | 26 | 2 |

**Table 3.** Results from searchlight RSA for the sensorimotor model. Table of coordinates and significance voxel-level peak values (p) in each activation cluster that was correlated with the sensorimotor semantic model.

| Regions | Cluster Extent | Voxel-level *P* | Pseudo *T* | Coordinates |  |  |
| --- | --- | --- | --- | --- | --- | --- |
|  |  |  |  | *x* | *y* | *z* |
| Left Inferior Temporal | 694 | 0.016 | 5.44 | -51 | -46 | -12 |
| Left fusiform |  | 0.016 | 4.45 | -39 | -46 | -24 |
| Left Mid Temporal |  | 0.016 | 3.60 | -48 | -40 | 4 |
| Left Occipital | 312 | 0.017 | 4.27 | -33 | -70 | 40 |
| Left Angular |  | 0.026 | 2.97 | -30 | -55 | -31 |
| Left Inferior Frontal Triangularis | 153 | 0.016 | 4.13 | -48 | 26 | 21 |
| Left Inferior Frontal Opercularis |  | 0.017 | 4.03 | -48 | 10 | 29 |
| Left Precentral |  | 0.017 | 4.52 | -48 | 14 | 31 |
| Left Mid Frontal | 45 | 0.027 | 4.05 | -33 | 23 | 48 |
| Right Middle Temporal | 196 | 0.018 | 4.23 | 51 | -64 | 10 |
| Right Middle Temporal |  | 0.018 | 4.20 | 45 | -73 | 21 |

**Table 4.**
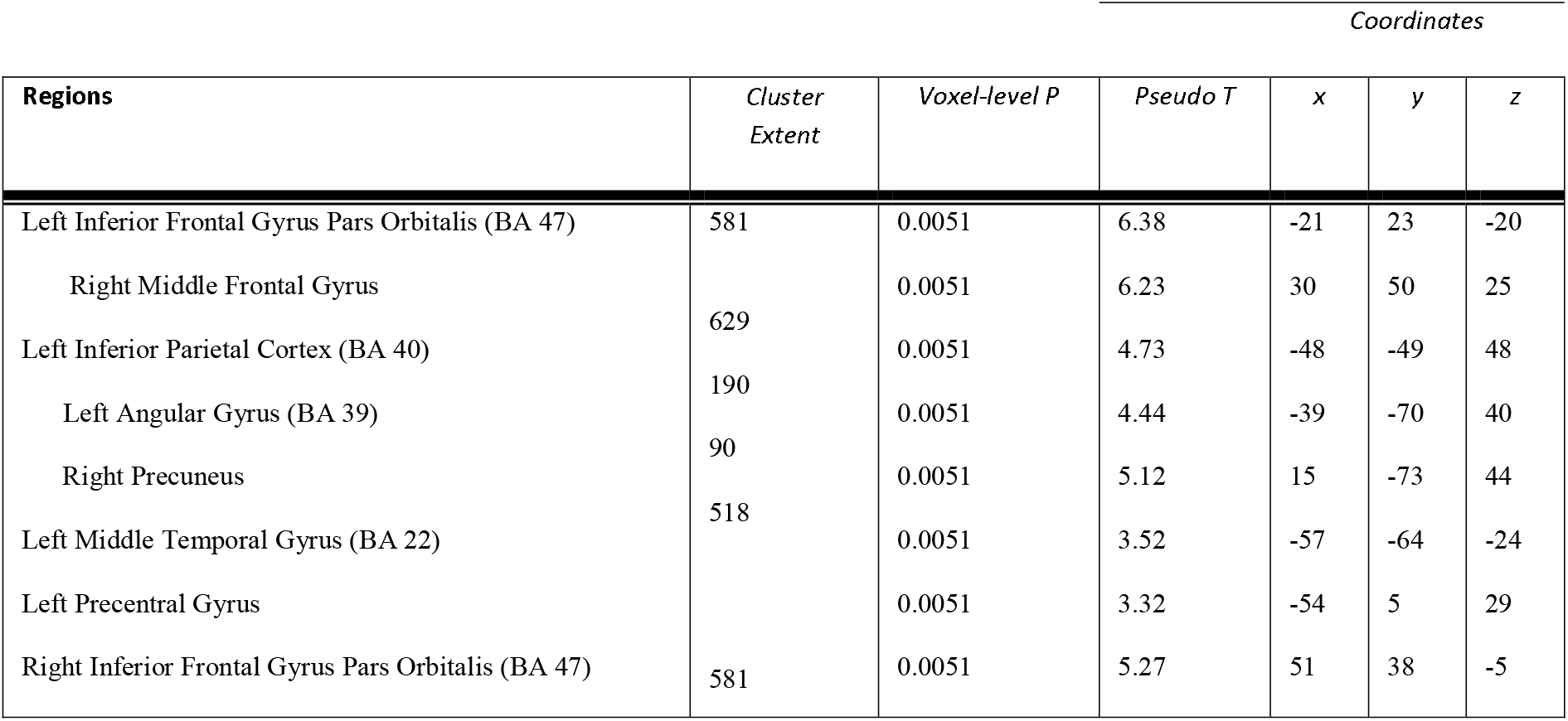
Results from searchlight RSA for the distributional model. Table of coordinates and significance voxel-level peak values (p) in each activation cluster that was correlated with the distributional semantic model.

### Results from ROI analyses

Numerically, the integrative model performed best (i.e., produced the most significant correlations across subjects) in pITG-MTG and LIFG (BA 44-45) with additional significant effects in left angular gyrus, precentral, anterior temporal cortex and fusiform gyrus (see Table 5). Likewise, the grounded sensorimotor model revealed most significant effects across subjects in pITG-MTG, LIFG BA44-45, and angular gyrus (Table 6). The distributional LSA model led to significant effects in more focused regions, such as the LIFG, and anterior ITG (Table 7).

**Table 5.** ROI results for the integrated model. Table of correlation values (*r*) and significance values (*P*) between the brain activity patterns in ROIs and the model combining sensorimotor and distributional information for all words. Correlations values which survive FDR correction for multiple comparisons and the model are indicated by asterisk (*).

| ROI | p-val | sig |
| --- | --- | --- |
| LIFG BA44 | 0.000319 | ** |
| LIFG BA45 | 0.000127 | ** |
| LIFG BA47 | 0.001363 | ** |
| PCG | 0.003726 | * |
| TP | 0.600114 |  |
| aITG | 0.032518 | * |
| aMTG | 0.157277 |  |
| aSTG | 0.7774 |  |
| pITG | 9.07E-05 | ** |
| pMTG | 0.006764 | * |
| pSTG | 0.172342 |  |
| FuG | 0.001208 | ** |
| SMG | 0.016335 | * |
| AG | 0.012711 | * |

**Table 6.** ROI results for the sensorimotor model. Table of correlation values (*r*) and significance values (*P*) between the brain activity patterns in ROIs and the sensorimotor model for all words. Correlations values which survive FDR correction for multiple comparisons and the model are indicated by asterisk (*).

| ROI | p-value |  |
| --- | --- | --- |
| LIFG BA44 | 0.000175 | ** |
| LIFG BA45 | 0.001363 | ** |
| LIFG BA47 | 0.037435 |  |
| PCG | 0.006764 | * |
| TP | 0.699477 |  |
| aITG | 0.059323 |  |
| aMTG | 0.196536 |  |
| aSTG | 0.811739 |  |
| pITG | 5.33E-05 | ** |
| pMTG | 0.001726 | ** |
| pSTG | 0.365694 |  |
| FuG | 0.001726 | ** |
| SMG | 0.020763 | * |
| AG | 0.006147 | * |

**Table 7.** ROI results for the distributional model. Table of correlation values (*r*) and significance values (*P*) between the brain activity patterns in ROIs and the LSA model for all words. Correlations values which survive FDR correction for multiple comparisons and the model are indicated by asterisk (*).

| ROI | p-value | sig |
| --- | --- | --- |
| LIFG BA44 | 0.067067 |  |
| LIFG BA45 | 0.020763 |  |
| LIFG BA47 | 7.63E-05 | ** |
| PCG | 0.052271 |  |
| TP | 0.032518 |  |
| aITG | 0.003355 | * |
| aMTG | 0.084762 |  |
| aSTG | 0.055713 |  |
| pITG | 0.730041 |  |
| pMTG | 0.123375 |  |
| pSTG | 0.006764 | * |
| FuG | 0.446606 |  |
| SMG | 0.024221 |  |
| AG | 0.071209 |  |

Among these regions, comparisons of within-category similarity values for each semantic type with the between-category similarity values revealed better category discriminability in the patterns from left posterior inferior and middle temporal cortex and parietal regions, which best correlated with the integrative model (see Table 8). Category-specific regions in left precentral cortex and posterior section of the LIFG (BA 44), supporting action semantics, reflected the action-related information common to action words and action-related tool and food words. The patterns from left anterior LIFG (BA 47) and anterior temporal cortex did not show any category discrimination, in coherence with their expected role in general semantic processes.

**Table 8.** Results for category discriminability per ROI. Table of significance values (*P*) between the within- and between-category values of the brain activity patterns in each ROI. Asterisk indicate values which survive Bonferroni correction for multiple comparisons (*).

| ROI | p-value | sig |
| --- | --- | --- |
| LIFG<br>BA44 | 0.187705 |  |
| LIFG<br>BA45 | 0.318988 |  |
| LIFG<br>BA47 | 0.502391 |  |
| PCG | 0.277525 |  |
| TP | 0.5 |  |
| aITG | 0.249044 |  |
| aMTG | 0.052613 |  |
| aSTG | 0.45113 |  |
| pITG | 0.003898 | ** |
| pMTG | 0.005547 | * |
| pSTG | 0.017238 | * |
| FuG | 0.010379 | * |
| SMG | 0.007945 | * |
| AG | 0.014183 | * |

### Results from statistical model comparisons respectively reveal distributional- and sensorimotor-specific similarity in LIFG (BA 47) and pITG

Results from direct statistical model comparisons teasing apart the representations of distributional and sensorimotor semantics in the abovementioned ROIs showed significant differences between them in pITG and in LIFG BA 47, respectively (Fig. 3.B). The sensorimotor and integrative models performed comparably.

### Category-specific representational similarity patterns

Additional ROI analyses were performed to compare semantic similarities between the large lexicosemantic categories of action related verbs and object related nouns. A statistical analysis of pooled values from modality-preferential frontal and posterior areas (see Methods) indicated a differential mapping of lexicosemantic categories (ANOVA design: Word Category (action verbs vs. object nouns) x Region (frontal vs. temporo-occipital)). For the integrated sensorimotor and distributional semantics, results revealed a significant cross-over interaction between the factors Region and Word Category (F [1, 22] = 7.45, p = 0.01) (see Fig. 4.A). This effect is an indication of a double dissociation in the semantic similarity mappings of action and object words onto focal motor and visual regions. Action words triggered significantly stronger semantic mapping effects in inferior frontal and motor regions (LIFG BA44-45, a region in PCG non adjacent to inferior motor cortex and BA44) as compared with object nouns (p=0.036). Conversely, in temporo-parietal regions -including integration areas of TP (e.g., Patterson et al., 2007), FuG (e.g., Mion et al., 2014), and AG (e.g., Binder and Desai, 2011), the opposite was found (p=0.04). The sensorimotor model alone led to a comparable interaction effect (F [1, 22] = 7.85, p = 0.012). The distributional model did not reveal any semantic mapping differences across word categories.

**Figure 4.**
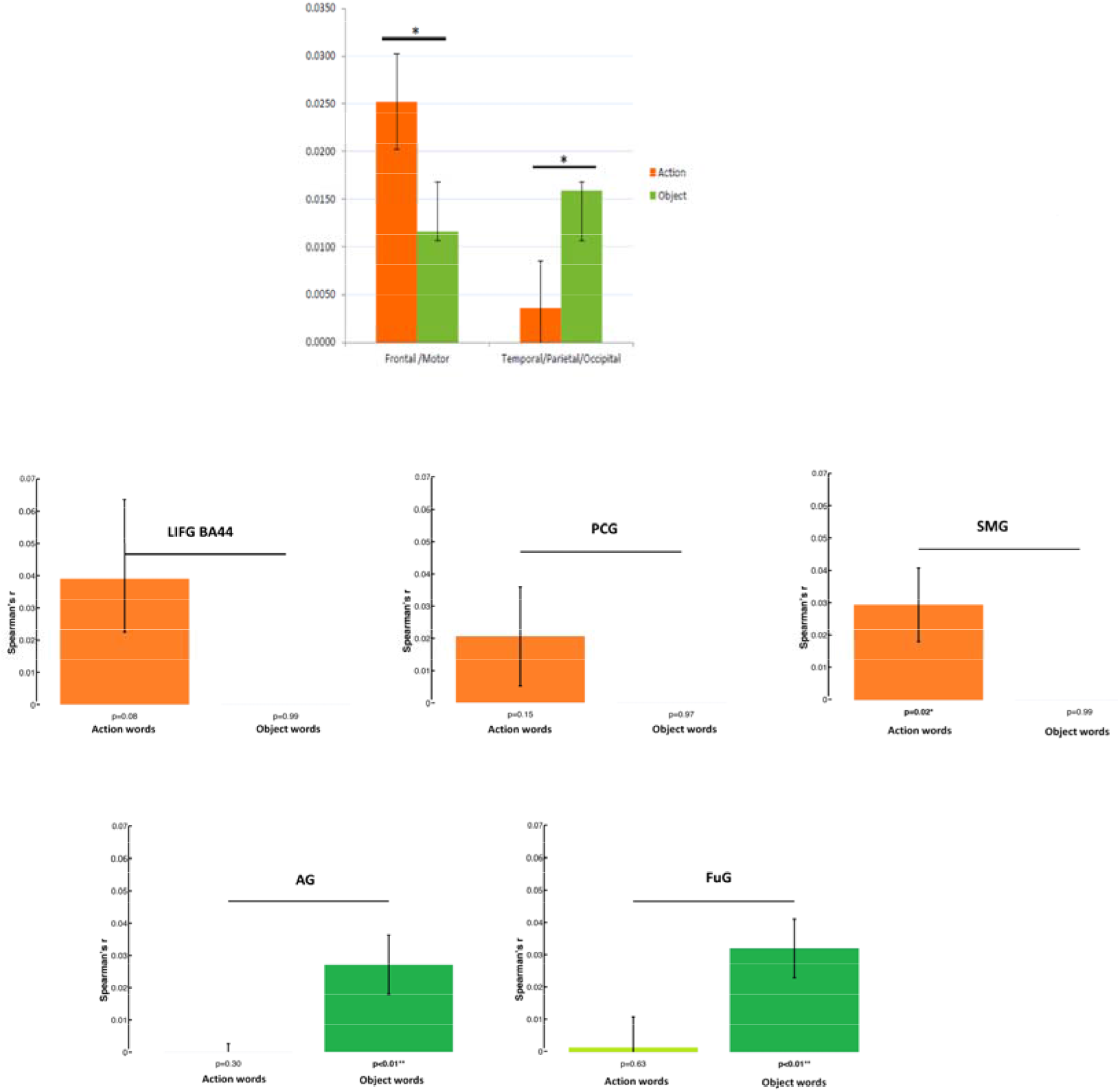
**A**: Results from statistical analyses showing differential mapping of lexicosemantic categories obtained (ANOVA design: Word Category (action verbs vs. object nouns) x Region (frontal vs. temporo-occipital)), with a significant cross-over interaction between the factors Region and Word Category (F [1, 22] = 7.45, p = 0.01). This effect was obtained by employing the integrated semantic vector model, and indicated a double dissociation in the semantic similarity mappings of action and object words onto focal motor and visual regions. A comparable interaction effect was also found when the sensorimotor model alone was applied (F [1, 22] = 7.85, p = 0.012). The distributional model did not reveal any semantic mapping differences across word categories. **B.** Category-specific results for object- and action-related concepts from model comparisons. The bar graphs depict the averaged correlations between brain activity in specific brain regions and the models based on motor properties (top panels) and visual properties (bottom panel) for the sub-spaces of action and object words. Significant differences are indicated by the horizontal bars in black after FDR correction across models (FDR=0.05). Top panel: in LIFG (BA 44), PCG and SMG, the model based on motor properties was correlated more strongly with the similarity patterns specific to action words as compared to the ones specific to object words. Bottom panel: in the FuG and AG, the visual properties correlated with the similarity patterns specific to object words more strongly than with the ones specific to action words.

### Results from motor and visual model comparisons reveal a double dissociation of action and object words in motor and temporo-parietal regions

Whilst directly contrasting the sensorimotor and the distributional sub-models for action and object words separately did not show any category-specific differences in any regions, the direct statistical comparisons between the models of the visual and motor properties for action and object words revealed a clear differentiation between the underlying representations in the respective action and perception systems (see Fig. 4.B).

These significantly distinct patterns (FDR=0.05) clearly pointed to a double dissociation between the representation of action and object words based on their respective sensorimotor attributes. These results confirm category-specificity in model-performance differences in motor and higher-order visual regions supporting the comprehension of action and object words.

## Discussion

In the present study, we investigated the relative contribution of distributional and sensorimotor semantics to the neural encoding of conceptual meaning. We found that, already on the level of purely cognitive linguistic information coded by the models, the integrative model combining sensorimotor and distributional information yielded better discrimination among conceptual categories than each of the two models taken in isolation. On the neural level, combined whole-brain RSA searchlights and ROIs analyses revealed a complex mapping of semantic similarity in distributed sets of high-level frontal and temporo-parietal regions in multimodal association cortex, also reaching primary motor and perceptual (visual) cortex for the encoding of specific action-related and object-related attributes. In this network, the patterns in left inferior temporal regions which more strongly correlated with the integrative model also showed best conceptual category discrimination. In contrast, the response patterns in left precentral regions and the posterior section of the LIFG (BA 44) correlated with the action-related information across categories (e.g., action concepts and action-related object concepts such as tools and foods). No category discrimination emerged from the patterns in left anterior LIFG (BA 47) and anterior temporal cortex, which were most sensitive to distributional information.

The results support theories positing sensorimotor simulations as being necessary for a precise discrimination and richer representation of concepts (Louwerse, 2008; Barsalou et al., 2008), and the integration of sensorimotor information with distributional statistics as being essential for the comprehension of the symbols used to name them (Barsalou, 2017; Pulvermuller, 2018).

### The need for an integrative semantic model

Previous behavioural work demonstrated that an extended range of 65 experiential attributes of concepts (e.g., sensorimotor, spatial, temporal, affective, social and cognitive components), as motivated by well-known neurobiological correlates, could be used to differentiate single items from several conceptual categories based on their relative semantic similarities (Binder et al., 2016). Interestingly, such brain-based experiential vectors were more accurate in capturing semantic similarities within the members of the same category than a 300-dimensional LSA vector over a large text corpus (see Binder et al., 2016 for discussion). In analogy with such earlier behavioural findings, the present data reveal that our semantic rating model leads to clearest differentiations between the similarity patterns of action- and object-related concepts relative to the distributional approach, but the integrative model combining both metrics reaches the most robust category discrimination. This confirms the theoretical assumption that distributional information resulting from the computational manipulation of symbols alone is insufficient for making those symbols interpretable. As postulated by the well-known Searlean Chinese room example, a computer could manipulate strings of Chinese symbols so well to perform as convincingly as a Chinese speaker would, even passing the Turing test, yet it would not understand their meaning (Searle, 1980; Harnad, 1990). Understanding linguistic symbols involves accessing the experiential attributes that link their arbitrary symbolic forms to their referents in the world. The grounding of arbitrary symbolic word forms in action and perception systems, where the information about referential meanings of at least an initial set of acquired concepts is stored consistently in terms of their perceptual/sensory and motor attributes (“grounding kernel”: Cangelosi and Harnad 2001), is thus essential for building the referential links between the arbitrary and symbolic word forms and the actions, objects and entities in the external world they are used to talk about (Harnad 1990, 2012; Pulvermüller 2013; Pulvermüller 2018). In the present data, the integrative model reflected the relative strength of the correlation for distributional and sensorimotor properties of words within and between semantic word categories and thus the ease with which statistical co-occurrence and sensorimotor information can be combined, thus enabling coherent representations of the words’ thematic structure based on its components. The present data are consistent with the view that both linguistic context and the action and object context, in which words and concepts are experienced since the early stages of language acquisition, critically contribute to coherent representations of lexical meaning (Wittgenstein, 1953).

Overall, our results from direct measure of metabolic signals during word comprehension support the theoretical assumption that the combinatorial knowledge of statistical regularities with which linguistic symbols co-occur in intralinguistic contexts (e.g., texts and media) does not give raise to the corresponding semantic representations (Harnad, 1990; Cangelosi and Harnad, 2001; Searle, 1980; Vigliocco et al. 2004; Andrews et al., 2014). Rather, a *condition sine qua non* for a complete picture of the semantic similarity space is the grounding of these symbols in the motor, perceptual, and affective properties of their referents in the “world”, as mediated by dedicated action and sensory systems (Barsalou, 2008; 2017).

Earlier work demonstrated that conceptual similarity, as evaluated by classical Latent Semantic Analysis of distributional statistics, could be mapped across the language regions, and especially in left inferior frontal gyrus (Carota et al., 2017). Recent findings brought further evidence for neuroanatomically separable, but representationally interdependent left inferior frontal and posterior middle/inferior temporal correlates of distributional and hierarchical/taxonomic similarity structure of concepts, respectively (Carota et al., 2021). In both studies, the LIFG emerged as a prominent site for distributional semantics, a result that is confirmed by the current data. Furthermore, both studies suggested that the similarity structure of words such as action concepts, food and tool object concepts, could be encoded based on their shared action-related properties. However, differences in the semantic similarity within and between action and object concepts based on sensorimotor and distributional information were not directly compared and their neural discriminability could not be uncovered in our previous work.

The present data contribute the state-of-art literature showing that the combination of sensorimotor and distributional semantic information yields best category discrimination of the object- and action-related words in the representational semantic space, as becomes especially manifest in temporoparietal regions. This finding extends the result from earlier work (Harnad, 1990; Cangelosi and Harnad, 2001; Vincent-Lamarre et al., 2016; Anderson et al., 2019), suggesting that integrative models of sensorimotor and distributional representations lead to a better approximation of human semantic similarity judgments of grounded experiential features as compared to the use of each estimate taken separately.

### Whole brain semantic similarity mapping

As a first finding of the present study, we show that the integrative model combining grounded sensorimotor and distributional statistics described the brain’s multidimensional semantic space discriminating among semantic categories. A comparable performance was seen for the model coding for sensorimotor semantic properties. A second major result showed that both the integrative and the sensorimotor model were more distinct from the distributional model in pITG, whereas the distributional semantic vector selectively correlated with the patterns in LIFG BA 47.

As a further result of importance, according to the integrative model and grounded sensorimotor models, the semantic spaces of action related verbs and object related nouns were better mapped in specific modality-preferential frontal (motor) and temporo-parietal (visual) areas, respectively. Furthermore, we also observed that only if the pertinent set of action-related and visually-related properties was selected for model comparison, a clear double dissociation became manifest in the spatial distribution of their respective representations in modality-preferential premotor, and higher-level visual regions in posterior temporal and parietal cortex. Taken together, these results suggest that only an increase in perceptual simulation due to the integration of sensorimotor experiential information with distributional knowledge allows for more precise and complete conceptual representations.

### Hierarchical architecture of the representational semantic systems: category-general semantic similarity mapping in higher-level association cortex

The present results highlight a hierarchical architecture of the representational systems encoding lexical meaning, according to which distributional and grounded experiential semantic properties of linguistic symbols are represented in different sets of modality-general and modality-specific regions in motor and sensory cortex. Our data are consistent with models positing the co-presence of multimodal semantic properties in several higher-order convergence zones holding conjunctive representations derived from multiple low-level sensory and motor representations of concepts (Binder and Desai, 2011; Damasio, 1989a, 1989b; Meyer and Damasio, 2009). Indeed, we found that the LIFG (BA 47) and the pITG best reflected similarity in distributional and experiential properties of the stimulus words, respectively, whereas the AG coded for both types of semantic information. This is interestingly in line with results from connectivity analyses showing that a cortical network comprising these same regions in left inferior frontal (BA 47), posterior inferior temporal, and inferior parietal cortex supports semantic processing functions specifically (e.g., Xiang et al., 2010). In particular, posterior inferior temporal cortex is known to store lexical information about conceptual features in memory (Mitchell et al., 2008; Tyler et al., 2013; Fairhall and Caramazza, 2013; Devereux et al., 2013; Carlson et al., 2014; Hagoort, 2019; Mitchell and Cusack, 2015; Ghio et al., 2016; Borghesani et al., 2016; Coutanche et al., 2016), thus supporting a semantic route to reading (e.g., Price, 2000). The AG encodes cross-modally integrated representations of semantic knowledge (e.g., Fernandino et al., 2015; 2016), as well as category-general representations of statistical knowledge, as our present results show. This area may thus be a multimodal association region binding together different formats and modalities of semantic representations to enable the comprehension of complex conceptual information. For instance, earlier evidence suggests that the AG supports the representation of complex concepts (e.g., plaid jacket), based on simple conceptual constituents (e.g., jacket and plaid) (Price et al., 2015). The semantically coherent representation of these conceptual combinations presupposes the multimodal integration of cross-modal attributes in this well-established semantic “hub” area (Binder et al., 2009; Fernandino et al., 2015; Binder and Desai, 2011). The interplay of distributional and experiential information in the AG may be an important prerequisite for the representation of the thematic structure of the words, a proposal that fits well existing neuropsychological evidence for semantic associative naming errors in aphasic patients exhibiting lesions to this region (Schwartz et al., 2011), as well as recent neuroimaging results (Xu et al., 2016; Carota et al., 2020).

Neuronal populations coding for representations of lexical-semantic similarities in the posterior temporal and parietal (AG) memory circuit (e.g., Fuster, 1997; Hagoort, 2016; 2019) may communicate via long-distance connections with population codes in inferior frontal regions, especially in the LIFG (BA 47), as shown across imaging modalities (e.g., Lau et al. 2008; Xiang et al., 2010; Hagoort, 2005; 2015). Strengthened activity in this network may arise as a result of spread of activation from the LIFG. In turn, the more anterior/ventral aspect of the left inferior frontal cortex is critical for unification of semantic information into linguistic structures (e.g., Hagoort, 2016), a process which operates on combinatorial semantic information and which may require access to the representation of statistical knowledge, such as the one captured by our distributional model. The present results fit this picture from previous literature well, stressing the importance of the LIFG (BA 47) in semantic processing (e.g., Poldrack et al., 1999; Thomspon-Schill et al., 1997), particularly for the encoding of distributional semantic similarity, a finding confirming earlier results (Carota et al., 2017; 2020).

Interestingly, neurophysiological results obtained using methods with a more precise temporal resolution than what fMRI provides have pointed out that the ITG-MTG may give input to the critical language regions, such as inferior frontal cortex (BA 45-47), anterior temporal and inferior parietal cortex, particularly the AG (e.g., Lau et al., 2008), thus enabling access to activated lexical-semantic representations. In the present work, anterior temporal regions showed a relatively stronger correlation of activity patterns with the distributional semantic model, which however did not reach a significant difference with the other two models, and did not stand out from the other language regions. In the context of the present study, it is possible that the absence of explicit semantic tasks like the silent reading task we employed might have led to a reduced activation of this region (Visser et al., 2010). Furthermore, the anterior temporal regions may be sensitive to more complex sets of feature conjunctions (e.g., Tyler et al., 2013) than the ones incorporated in our models. In conclusion, the present study confirms a role of the anterior temporal regions in the encoding of semantic similarity, but failed to confirm this region as a possible apex for all forms of semantic knowledge to be brought together, as expected on the basis of “Hub-and-spokes” model (Patterson et al., 2007; Lambon-Ralph, 2017; Hoffmann et al., 2018), highlighting its contribution within a widespread network of distributed conceptual representations.

### Category-specific representations of action-related and visually-related words

Consistent with current theories (e.g., Simmons and Barsalou, 2003), the present data reveal a hierarchy spanning from this set of higher-level cortical regions to intermediate levels of representations reaching modality-preferential regions. In particular, the data show a representational gradient in their underlying semantic space, from category-general multimodal representations in the network of frontal and temporo-parietal regions just mentioned, to category-specific representations of specific motor and visual attributes defining, respectively, action and object semantic word types, which also engaged the corresponding action and perception systems.

A major finding of the present study was that a sub-network of inferior parietal (AG) and fusiform (FuG) regions reflected object-specific representations of visually-related semantic similarity. Previous fMRI studies have shown that the preferential response of different portions of the FuG to different object categories, for example animals and tools as employed here (Warrington & Shallice, 1984; Chao, et al., 1999; Martin, 2007; Moss et al., 2005; Tyler et al., 2013; Forseth et al., 2018). Such categorical differentiation has been proposed to arise from higher and lower levels of shared features across the members of different categories of objects, giving origin to category structure (Tyler et al., 2013). It is interesting that, in the present data, the AG may become co-activated with an object-specific FuG region in the ventral visual pathway to code for an intermediate and more specialised level of sensory representation than the category-general information manipulated when co-activated with the fronto-temporal network discussed above.

In the present data, when the sub-classes of action and object concepts were examined separately, no category-specific differences were observed in the respective semantic sub-space of distributional and sensorimotor information, as expected on the basis of the theoretical assumption that word semantic categories require increased access to specific distinctive sensorimotor properties of specific classes of words. One may argue that the action-related nature of our test object words might have led to the lack of differential activity in the corresponding neural patterns relative to action words. However, despite the inclusion of action-related object words, the results from the comparisons of the models coding for specific visual and motor properties clearly rule out such possibility, demonstrating their critical contribution to the neurocognitive distinctiveness, and distinction, of action and object words. Our present findings thus point to a genuinely sensorimotor semantic origin of category-specific representations: classifying an object and the word used to speak about it as a member of a category may require simple conjunction of sensorimotor attributes, which confines activity to category-specific temporo-parietal regions. In the current data, it was noteworthy that specialized sub-networks encode distinctive information supporting the disambiguation and identification of specific categories of words and concepts.

The semantic sub-space in left motor, inferior frontal (LIFG BA 44), motor, and the SMG was, in turn, well mapped by the model coding for rated action-related properties of the words, consistent with earlier findings on the role of these regions in action semantics (e.g., Damasio and Tranel, 1993; Kemmerer and Tranel, 2003; Hillis et al., 2004, 2006; Rueschmeyer et al., 2010; Grossman et al., 2008; Kemmerer et al., 2012; Davey et al., 2015; Vukovic and Shtyrov, 2019; Dreyer et al., 2020).

## Conclusions

We showed that distributional and sensorimotor semantics is reflected by graded activation of neuronal populations in partly distinct networks of frontal and temporo-parietal regions. This neurocognitive picture informs semantic theories suggesting that a representationally complete characterization of the complexity of lexical meaning cannot be based on distributional information alone, but requires the activation of sensorimotor knowledge. Our results from highly controlled stimuli using a simple listening “non-task” support distributed models positing a task-sharing between modality independent and modality-specific regions in encoding and linking together semantic properties of words to form rich representations of lexical meaning in coherence with semantic word types. In the present data, the conjunction of a limited amount of specific sensorimotor properties, such as the dominant motor and visual attributes of action and object words, was sufficient for mapping the corresponding semantic spaces in modality-specific motor and visual cortex, thus corroborating the relevance of brain-based componential attributes of words and concepts (Binder et al., 2016; Fernandino e al., 2016; Pulvermüller et al., 1999; 2017). Future work is needed for determining to what extent the different sources of semantic information (e.g., distributional and sensorimotor properties of words, and their interplay, as employed here) contribute to the mapping of the representational content of language beyond single words (e.g., sentence, discourse, naturalistic contexts) in relevant brain networks.

